# Reconstructing the Gigabase Plant Genome of *Solanum pennellii* using Nanopore Sequencing

**DOI:** 10.1101/129148

**Authors:** Maximilian H.-W. Schmidt, Alxander Vogel, Alisandra K. Denton, Benjamin Istace, Alexandra Wormit, Henri van de Geest, Marie E. Bolger, Saleh Alseekh, Janina Maβ, Christian Pfaff, Ulrich Schurr, Roger Chetelat, Florian Maumus, Jean-Marc Aury, Alisdair R. Fernie, Dani Zamir, Anthony M. Bolger, Bjöern Usadel

**Author notes:** These authors contributed equally (Maximilian H.-W. Schmidt, Alxander Vogel, Alisandra K. Denton).

## Abstract

Recent updates in sequencing technology have made it possible to obtain Gigabases of sequence data from one single flowcell. Prior to this update, the nanopore sequencing technology was mainly used to analyze and assemble microbial samples^1-3^. Here, we describe the generation of a comprehensive nanopore sequencing dataset with a median fragment size of 11,979 bp for the wild tomato species *Solanum pennellii* featuring an estimated genome size of ca 1.0 to 1.1 Gbases. We describe its genome assembly to a contig N50 of 2.5 MB using a pipeline comprising a Canu^4^ pre-processing and a subsequent assembly using SMARTdenovo. We show that the obtained nanopore based *de novo* genome reconstruction is structurally highly similar to that of the reference *S. pennellii* LA716^5^ genome but has a high error rate caused mostly by deletions in homopolymers. After polishing the assembly with Illumina short read data we obtained an error rate of <0.02 % when assessed versus the same Illumina data. More importantly however we obtained a gene completeness of 96.53% which even slightly surpasses that of the reference *S. pennellii* genome^5^. Taken together our data indicate such long read sequencing data can be used to affordably sequence and assemble Gbase sized diploid plant genomes.

Raw data is available at http://www.plabipd.de/portal/solanum-pennellii and has been deposited as PRJEB19787.

## Results and Discussion

*Solanum pennellii* is a wild, green fruited tomato species native to Peru. It exhibits beneficial traits such as abiotic stress resistances^6,7^. The accession LA716 has been used to generate a panel of introgression^8^ and backcrossed introgression^9^ lines which have been used to identify several thousand QTL thus ideally complementing large scale genomic panel studies for tomato^10,11^. However, LA716 does not perform well in the field and carries the necrotic dwarf gene on chromosome 6 which reduces plant vigor when introduced into a *Solanum lycopersicum* background^12^. To overcome these problems, the accession LYC1722 was identified in a large panel of tomato accessions obtained from the IPK genebank in Gatersleben, Germany, as a self-compatible, phenotypically uniform biotype of *S. pennellii* that does not exhibit these negative traits of LA716. In addition, the metabolite content of LYC1722 is quite different from published values for LA716 with considerably higher levels of folial amino acids and lower levels of organic acids (supplemental Figure 1).

Given that, the Oxford Nanopore technology does not require a large capital investment, unlike other next generation sequencing methods, we used nanopore technology to sequence and assemble the genome of *S. pennellii* accession LYC1722. First, we generated about 39 Gigabases of 2x 300 bp Illumina reads. This data set revealed that this accession of *S. pennellii* has an estimated genome size of between 1 and 1.2 Gbases, similar to the estimate for the reference *S. pennellii* LA716. Further, this accession is relatively homozygous (Supplemental Figure 2) in line with its self-compatibility, a trait found in some southern *S. pennellii* populations, including LA716 and LA2963, and which contrasts with the strict self-incompatibility and high heterozygosity typical of this species as a whole^13^. Using the short read sequencing data to identify variants such as single nucleotide polymorphisms (SNPs) and small insertions and deletions (InDels) versus the *S. pennellii* LA716 reference revealed 6.2 million predicted variants where the highest variant rate was found on chromosomes 1 and 4 (Figure 1A, Supplemental Table 1). This indicated a relatively large difference between this accession and the reference, as in a large panel of cultivated tomatoes (*S. lycopersicon*) more than 2 million variants were only found in a few cases^14^. This is further evidence of the high level of diversity within *S. pennellii*^13^ and shows that LA716 and LYC1722 are relatively diverged populations.

**Figure 1.**
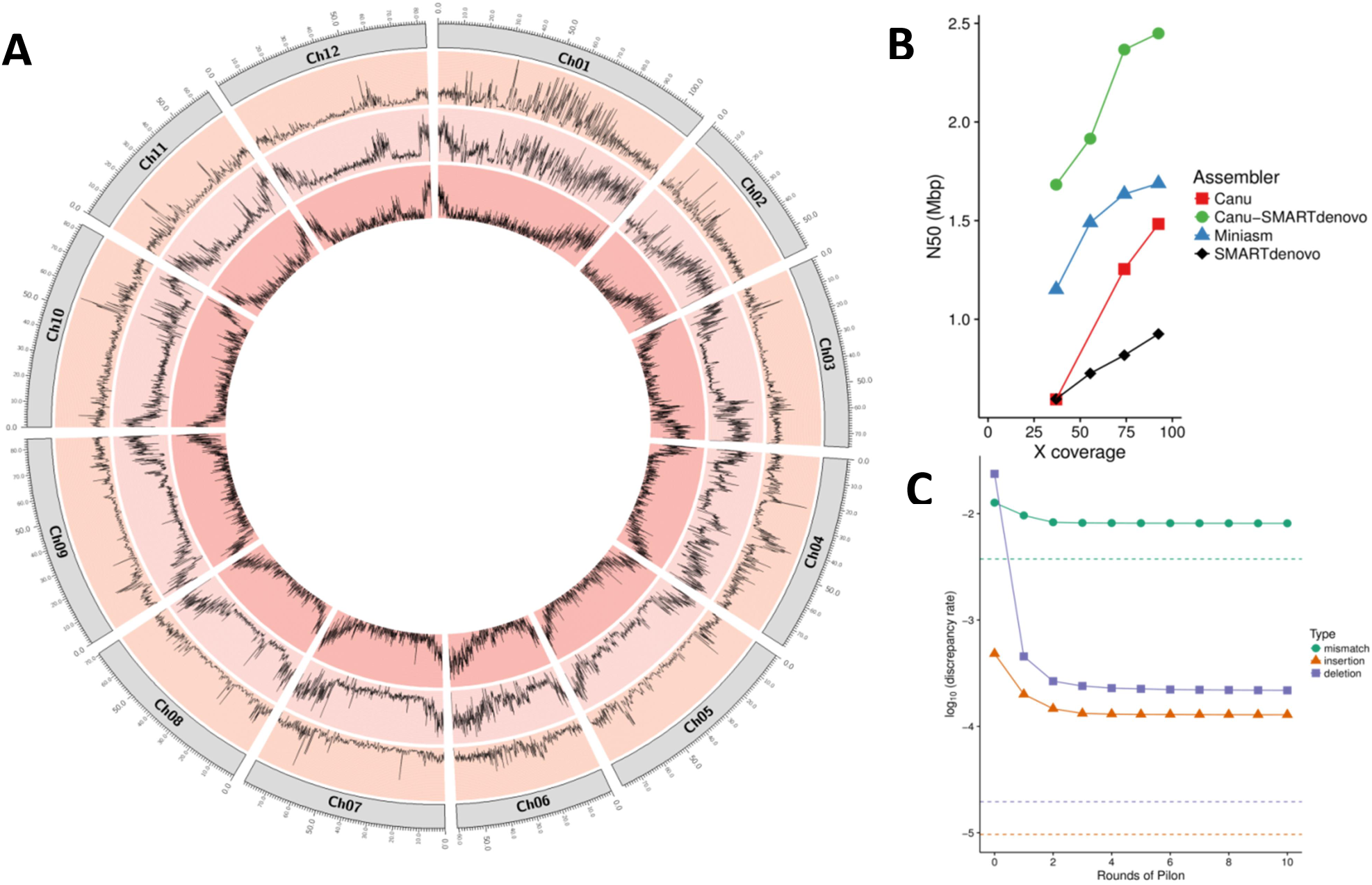
Characteristics of the LYC1722 genome and its assembly. A) Circos visualization of variant distribution between *S. pennellii* LYC1722 and *S. pennellii* LA716 Distribution of SNPs (outer layer) and InDels (middle layer) is compared to the gene density (inner layer) for each chromosome of *S. pennellii* LA716 based on generated Illumina data for *S. pennellii* LYC1722. B) The effect of randomly downsampling pass reads on the N50 produced by different assemblers C) Discrepancies between the assembly and the Illumina data over several rounds of Pilon correction. Dotted lines approximate expected discrepancy rates if Illumina data were mapped to a perfect reference.

We therefore sequenced the genome of this new self-compatible *S. pennellii* accession with Oxford Nanopore reads. Thirty-one flowcells yielded 131.6 Gbases of data in total, of which 110.96 Gbases (representing about 100 fold coverage), were classified as “passed filter”, by the Oxford Nanopore Metrichor 1.121 base caller. As shown in Supplemental Table 2, total yield per flowcell varied between 1.1 and 7.3 Gbases before and 0.96 and 6.02 Gbases after filtering and most data was obtained within the first 24h of sequencing (Supplemental Figure 3). The average Q-score was around 6.88 and 7.44 before and after filtering (Supplemental Figure 4). Re-aligning the reads revealed a typical read identity value of aligned bases of 80.97% (Supplemental Figure 5), in line with the obtained quality values shown (Supplemental Figure 6). However, these values are lower than observed in microbial data reflecting older pore data values^15,16^, which could be explained by the fact that the base caller was not trained for plants.

The average read length varied between 6463 (6625) and 14901 (15869) with library preparation optimizations before (and after) quality filtering (Figure 2, Supplementary Table 4). This is in part due to the preparation, but it was possible to routinely achieve 12.7 kb long reads when gel-based size selection was applied. The longest reads passing quality filtering was 153,099 bases long of with an alignment length of 132,365.

**Figure 2.**
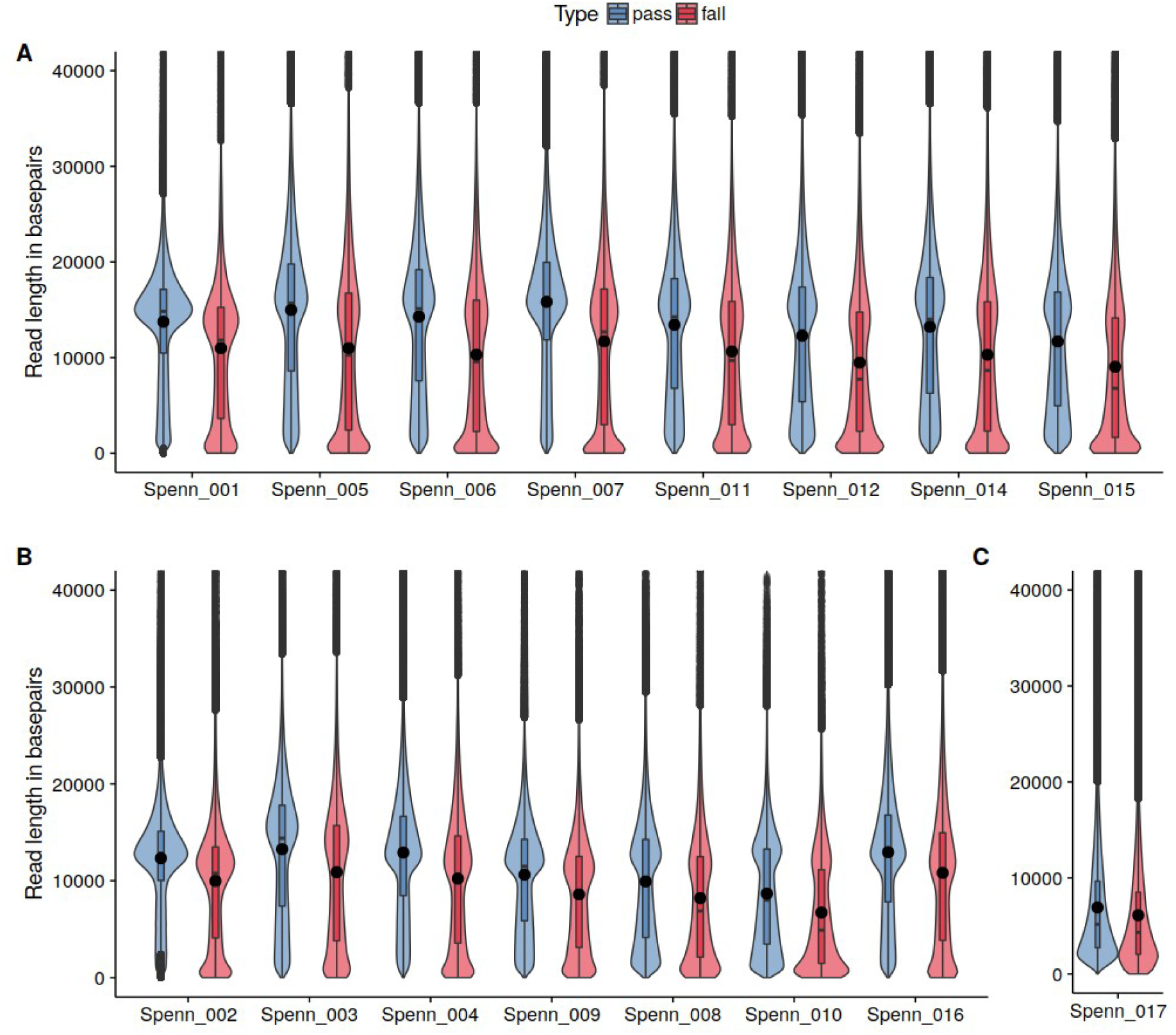
Violin-plots of read length per library for three different size selection protocols. Read-length distribution is shown for all 16 *S. pennellii* MinION libraries and the corresponding pass (blue) and failed (red) classified reads. Libraries are grouped by size selection protocol (A: 15 kb cutoff, B: 12 kb cut-off, C: 0.4x bead size-selection). Filled dots indicate mean read length.

Since plant genomes are known to be notoriously difficult to assembly due to their high fraction of repetitive DNA^17^, we wondered if the Oxford Nanopore long read data could be used to achieve a contiguous assembly reflecting the genome structure of *S. pennellii*. To assess this, we used the “passed filter” data comprising almost 100-fold estimated coverage and subsampled the data to 40, 60, and 80%. Each data set was assembled using miniasm^18^, Canu^4^ and SMARTdenovo which represent the state-of-the-art assemblers known to support Oxford Nanopore sequencing technology^19^. As can be seen in Figure 1B, the N50, a measure of the minimum contig length to cover 50% of the genome, was still rising for the subsampled data sets. This is in contrast to the Canu based assembly of the model plant *Arabidopsis* using long PacBio read data, as for this small genome Canu saturated at approximately 50 fold genome coverage at an N50 of more than 5 Mbases^4^. This is likely explained by the inclusion of more long reads relative to the genome size. To test this hypothesis, a 30 fold coverage subsample of the reads using different read length averages showed that SMARTdenovo alone produced N50 values of above 1 Mbase with only 30 fold coverage when average read length was above 20 kbases, whereas an average read length of less than 13kbases only yielded an N50 of about 0.2 Mbases (Supplemental Figure 7).This is in line with the result that when randomly subsampled to 40% of the data, all assemblers produced an N50 value above 0.5 Mbases. Assembling the genome with the hybrid assembler dbg2olc^20^ or using a very early version of the wtdbg assembler did not yield N50 values above a megabase for the full dataset and were thus not considered further (supplemental File A, B).

The largest N50 for the full dataset was obtained using miniasm with an N50 of 1.69 Mbases, versus 1.48 Mbases for Canu and 1.03 Mbases for SMARTdenovo (Table 1) after parameter tuning (supplemental File C, D). However to complete the assembly, Canu needed almost two orders of magnitude more CPU hours than miniasm or SMARTdenovo (Table 1). That said, miniasm does not correct assemblies^10^ at all, so the resulting “assembly” would need to be carefully post-processed. This is also reflected in the base error rate of the genome which was estimated by calling variants in this data using samtools^21^ using the Illumina read data. This revealed error rates of 2.66, 1.54 and 1.1, for miniasm, SMARTdenovo and Canu for regions covered by at least five reads (Supplemental File E). As variant calling may underestimate errors, we further calculated raw discrepancies between mapped Illumina reads and the assemblies with Qualimap. Qualimap counts differences between the assembly and individual Illumina reads thus providing a compound estimate for the sum of errors in the assembly, in the Illumina reads and for heterozygous positions. Here we obtained Qualimap discrepancy rates of 9.11, 4.22, and 3.74% for miniasm, SMARTdenovo and Canu, respectively (Supplemental File F). These values give an upper bound for the base error rates covered by Illumina reads. For Canu and SMARTdenovo most errors resulted from deletions. Most of the remaining errors were due to mismatches whereas errors resulting from insertions were more than 10 times less likely (See Supplemental File F). When aligning the assemblies versus the LA716 reference genome, we observed that all three assemblies were largely in agreement with the reference (Supplemental Figure 8).

**Table 1.** Assembly statistics and run-time statistics by assembly and post-processing (see also supplemental File for additional polishing data)

|  |  | k cpu<br>hours | Memory<br>(GB) | N50 | L50 | Total<br>size | Largest<br>contig | Total<br>contigs | Illumina<br>mapping<br>rate (%) | Qualimap<br>Discrepancy<br>rate | %<br>complete<br>BUSCO |
| --- | --- | --- | --- | --- | --- | --- | --- | --- | --- | --- | --- |
| Canu | Raw | 80.42 | 199.87 | 1.48 | 169 | 922.94 | 9.63 | 2010 | 98.52 | 3.74 | 26.46 |
| SMARTdenovo |  | 0.72 | 55.60 | 1.03 | 271 | 929.99 | 5.68 | 1901 | 98.65 | 4.22 | 26.74 |
| Miniasm |  | 1.86 | 51.93 | 1.69 | 158 | 956.29 | 9.28 | 2704 | 95.53 | 9.11 | 0.21 |
| Canu-SMARTdenovo |  | 10.68 | 131.32 | 2.45 | 106 | 889.92 | 12.32 | 899 | 98.73 | 3.68 | 29.1 |
| Canu | Pilon polished 5x | - | - | 1.55 | 169 | 961.83 | 10.01 | 2010 | 98.95 | 0.82 | 96.46 |
| SMARTdenovo |  | - | - | 1.06 | 270 | 955.31 | 5.84 | 1901 | 98.99 | 0.91 | 96.11 |
| Miniasm |  | - | - | 1.75 | 156 | 977.78 | 9.49 | 2704 | 98.24 | 2.48 | 85.69 |
| Canu-SMARTdenovo |  | - | - | 2.52 | 106 | 915.60 | 12.72 | 899 | 98.98 | 0.85 | 96.46 |

All sequence length in Mbp

In order to test the functional completeness of the genome we used BUSCO^22^ that tries to find conserved genes in the genome. Here we obtained a gene completeness estimate of 0.21, 26.46 and 26.74% for miniasm, Canu and SMARTdenovo using BUSCO, respectively (Table 1, Supplemental File G). This suggests that as anticipated all three assemblies, whilst being structurally mostly correct can only be considered pre-drafts and should not be regarded as useful for gene definition likely due to their high InDel rate. Given the high error rates identified in the assemblies, we suspected that these might also negatively impact the genome assemblies. Therefore we used Canu to pre-correct the original reads and assembled the resulting data using SMARTdenovo, as SMARTdenovo alone yielded the best BUSCO score. Indeed, we observed that the resulting assembly had a superior N50 of 2.45 Mbases in just 899 contigs. The largest contig obtained was 12.32 Mbases long and the estimated error rate for regions covered by at least 5 Illumina reads was 1.2 % whereas the Qualimap discrepancy rate was 3.68 % (Supplemental File E and F). Concomitantly the BUSCO completeness rate rose to 29.1% for this two tool assembly (Table 1).

The above-listed values were, however, still far from sufficient for a genome assembly. We therefore post-processed the assemblies using the Oxford Nanopore based assembly corrector racon^23^ and/or the Illumina based polishing tool Pilon^24^. As can be seen in Figure 1C (and the Supplemental File E,F), the error rate dropped consistently for each application of Pilon for the Canu assembly up to the 5^th^ round of error correction, where the raw mismatch rate approached that expected for the Illumina reads (0.83 % vs an expected 0.37%). That said, slight reductions continued through the 10^th^ round of Pilon. As stated above, Qualimap measures the sum of errors in the assembly and in the Illumina reads. The expected Illumina error rate based on the quality scores was 0.37%, accounting for almost half of the final discrepancy rate of 0.83%. Both expected Illumina errors and final discrepancy rate after polishing were dominated by mismatches. Furthermore, heterozygous positions contribute to this discrepancy rate as about 50% Illumina reads covering these will disagree with the assembly (Supplemental File E). More conservatively, when calling homozygous variants using the same Illumina data, we could identify fewer than 90,000 homozygous variants in 840 MB of genomic regions in the polished Canu assembly covered by at least five reads representing an error rate approaching 0.01%. Similarly, for the Canu-SMARTdenovo assembly a discrepancy rate of 0.86% and an error rate for homozygous variants of <0.02% was found (Supplemental File E).

The BUSCO gene completeness peaked at 96.53 % for the Canu-SMARTdenovo assembly after 4 rounds of Pilon, with further polishing resulting in small, stochastic, loses of up to 3 (0.2%) BUSCOs. This peak BUSCO completeness was slightly better than that of the reference genome for LA716 at 96.32 % (3 fewer BUSCOs). Using racon^23^, a corrector making use of the original Oxford Nanopore based data, the Qualimap discrepancy rate could be decreased to about 3.6% for the Canu and Canu/SMARTdenovo assemblies (Supplemental File E,F).

When comparing LYC1722 phenotypically to accessions from the Rick center we found the very similar self-compatible accession LA2963 and wondered if these were related. We therefore resequenced one LA2963 individual using Illumina technology at low coverage and aligned the read data to the LYC1722 genome. When only including regions (ca 600MBases) covered by at least two reads of high mapping quality, we were able to find only about 200k homozygous variants (supplemental File H) which is much lower than the remaining heterozygosity of LYC1722 represented by more than 500k heterozygous positions in the genome (supplemental file H). Thus it is quite possible that these two samples represent the same original accession, i.e. LA2963.

In conclusion, we demonstrated that it is possible to obtain functional and highly contiguous genome assemblies covering most of the gene space for Gbase sized plant genomes using nanpore based long read data. Given a bulk discount price of about $500 per flow cell, and a cost for $215 for library preparation per up to three flow cells, consumable costs for medium sized plant genomes would thus be estimated to be approximately $25000. Although additional major cost factors are he computation times, these are expected to fall, especially with more precise and eukaryote optimized basecallers. In addition further methodological improvements to get even higher average read length (cf. Figure 2) will decrease computational requirements and would also bring the coverage requirement down (Supplemental Figure 7).

## Methods

### Plant growth

*S. pennellii* LYC1722 seeds were surface sterilized in a 10 % hydrogen peroxide solution for 10 minutes, rinsed three times with sterile water and transferred to 0.8 % half strength Murashige and Skoog Gelrite plates supplemented with 1 % Sucrose and 10 μM Gibberellic acid. Seeds were incubated for 7 days under constant light at 22 °C in a CLF Percival mobile plant chamber at 110 μmol m^-2^ s^-1^ light intensity. Seedlings were transferred on soil and further cultivated in a greenhouse supplemented with artificial light to a light intensity of at least 200 μmol m^-2^ s^-1^ for 16 h a day.

*Solanum pennellii* LA2963 seeds were obtained from the C. M. Rick Tomato Genetics Resource Center and germinated the same way as *S. pennellii* LYC1722. Planteles were transplanted to Rockwool cubes irrigated with Hoagland media solution over a continuous dripping system in a Phytochamber with 400 μmol m^-2^ s^-1^ light intensity, 12 hours of light at 18°C and 70% humidity during light cycles and 15°C and 80% humidity during dark cycles.

### Long fragment enriched 1D R9.4 library preparation

In order to take advantage of the long read technology an optimized protocol for enrichment of DNA fragments of 12-20 kbp was developed based on Oxford Nanopore’s “1D gDNA selection for long reads” protocol. For compatibility with the R9.4 SpotON MIN106 flow cells the Ligation Sequencing Kit 1D (R9.4) was used (Oxford Nanopore Technologies, SQK-LSK108). For each library 20 μg of high-molecular weight DNA was sheared using a g-Tube (Covaris) in a total volume of 150 μl nuclease free water at 4500-6000 rpm depending on the desired fragment size. Enrichment for long fragments was achieved by BluePippin size selection (Sage Science). Approximately 35 μl per lane was run together with an S1 marker reference lane on a 0.75 % Agarose Cassette (Biozym) using the high pass protocol and a collection window of 12-80 kbp or 15-80 kbp. Upon completion of the elution, the sample was allowed to settle for at least 45 minutes to allow the long DNA fragments to dissociate from the elution well membrane. All subsequent bead clean-ups were performed with an equal volume of Agencourt AMPure XP beads (Beckman) with elongated bead binding and elution time of 15 minutes on a Hula Mixer (Grant) at 1 rpm/min. Bead binding was carried out at room temperature and elution at 37 °C. Subsequently up to 5 μg of DNA was used for NEBNext ^®^ FFPE DNA Repair (New England Biolabs) in a total volume of 155 μl including 16.3 μl NEBNext FFPE DNA Repair Buffer and 5 μl NEBNext FFPE DNA Repair Mix. The reaction was incubated for 15 minutes at 20 °C. To reduce DNA shearing during the following bead clean up, the sample was split in two 77.5 μl aliquots and eluted each in 50.5 μl nuclease free water. For NEBNext ® Ultra™II End Repair/dA-Tailing treatment (New England Biolabs) 100 μl of FFPE repaired DNA, together with 14.0 μl NEBNext Ultra II End Prep Reaction Buffer and 6 μl NEBNext Ultra II End Prep Enzyme Mix, were incubated for 30 minutes at 20 °C followed by 20 minutes at 65 °C and 4 °C until further processing. For purification the sample was split again into two aliquots of 60.0 μl and subjected to a bead clean up. 20 μl of Oxford Nanopore 1D Adapter Mix (1D AMX, Oxford Nanopore Technologies, Cat# SQK-LSK108) were ligated to 30 μl of end repaired and adenylated DNA with 50 μl NEB Blunt/TA Master Mix (New England Biolabs, Cat# M0367L) for 20 minutes at 25 °C. As the motor protein is already part of the adapter, beads were resuspended twice with Oxford Nanopore Adapter Bead Buffer (ABB, Oxford Nanopore Technologies). The final library was eluted in 13-37 μl of Oxford Nanopore Elution Buffer (ELB, Oxford Nanopore Technologies) depending on how many flow cells were run in parallel. The final sequencing library was kept on ice until sequencing, but time was kept as short as possible. An overview of intermediate DNA quantifications and clean-up recoveries can be found in Supplementary Table 3.

### Non size-selected library preparation

A total amount of 10 μg high-molecular weight DNA in 150 μl was used for g-Tube (Covaris) sheared at 4500 rpm. Directly after shearing 0.4x vol. Agencourt Ampure XP beads (Beckman) were added to the sample to deplete small fragments while following the bead clean up protocol with elongated bead binding and elution as described above. The bead size selected DNA was eluted in 133.7 μl nuclease free water. Based on Qubit dsDNA BR quantification 5 μg of DNA was subjected to the protocol described for long fragment enriched libraries from NEBNext FFPE DNA Repair to the adapter ligation. The ratio of Agencourt AMPure XP beads (Beckman) for the final bead clean-up of the ligation reaction was adjusted to 0.4x of the sample volume for repeated depletion of small fragments. The library was eluted in 25 μl for Qubit dsDNA BR (ThermoFisher Scientific) quantification and loading of two flow cells.

### MinION Sequencing

All sequencing runs were performed on MinION SpotON Flow Cells MK I (R9.4) (Oxford Nanopore Technologies, Cat# FLO-SPOTR9). Immediately before start of sequencing run and within five days of delivery, a Platform QC was performed to determine the number of active pores (Supplementary Table 2). Priming of the flow cell was performed by applying 800 μl priming buffer (500 μl Oxford Nanopore Running Buffer RBF and 500 μl nuclease free water) through the sample port. After 5 minute incubation at room temperature, 200 μl of priming buffer was loaded through the sample port with opened SpotON port. In parallel 12 μl of final library was mixed with 25.5 μl Library Loading Beads (Oxford Nanopore Technologies LLB) and 37.5 Running Buffer 1 (Oxford Nanopore Technologies RBF1). Directly after priming 75 μl of the prepared library was loaded through the SpotON port. Loading amounts of libraries quantified via Qubit dsDNA BR assay are given in Supplementary Table 2. The sequencing script “NC_48Hr_Sequencing_Run_FLO-MIN106_SQKLSK108” was used with disabled live base calling. Basecalling was performed upon completion of the sequencing run with Metrichor and the “1D Basecalling for FLO-MIN106 450 bps” workflow (v1.121).

### MiSeq Sequencing

High molecular weight DNA from one 2 month old plant of *S. pennellii* LYC1722 and one plant of LA2963 was extracted as described earlier^5^.

For *S. pennellii* LYC1722 2 μg of this DNA were sheared using a Diagenode Bioruptor Pico Sonicator using 5 cycles of 5 seconds sonication interchanging with 60 second breaks to yield fragmented DNA with a medium insert size of 550 base pairs. The fragmented DNA was then used to create an Illumina TruSeq PCR-free library according to the manufacturer’s instructions.

The sequencing library was quantified using the Perfecta NGS Quantification qPCR kit from Quanta Biosciences and sequenced four times on an Illumina MiSeq-Sequencer using three 600 cycle V3 and one 150 cycle V2 Sequencing Kits.

For *S. pennellii* LA2963 5 μg of high molecular weight DNA were sheared using a Diagenode Bioruptor using 8 cycles of 5 seconds sonication interchanging with 60 second breaks to yield fragmented DNA with a medium insert size of 350 base pairs. The fragmented DNA was then size selected from 200-500 base pairs using a Blue Pippin with Dye free 1.5% Agarose cartridges and Marker R2. Size Selected DNA was then purified using Beckman and Coulter Ampure XP beads in a sample to beads ratio of 1:1.6. To repair possible single strand nicks DNA was then treated with the New England Biolabs FFPE-repair-mix according to manufacturer’s instruction followed by another Ampure XP bead Clean-Up. DNA was then end-prepped and adenylated using the NEBNext Ultra II DNA Library Prep Kit according to manufacturer’s instructions. For Ligation of Sequencing Adapters 2.5 μL of Adpater 13 from the Illumina TruSeq PCR-free Kit was used together with the 30 μL of the NEBNext Ultra II Ligation Master Mix, 1 μL NEBNext Ligation Enhancer and 60 μL of the End Prep Reaction Mixture. These components were mixed and incubated at 20°C for 15 minutes before adding 3 μL nuclease-free water and incubating at 37°C for 15 minutes. Afterwards adapter ligated DNA was cleaned up with two consecutive bead clean-ups with a 1:1 ratio of sample and beads. The resulting Library was quantified using the NEBNext Library Quant Kit for Illumina and sequenced on an Illumina MiSeq-Sequencer using a 150 cycle V3 Sequencing Kit.

### Assembly

Base calling ‘pass’ reads were assembled with a variety of different tools to determine whether coverage was saturating. Then parameters and tool-combinations were further refined to obtain a handful of ‘top’ assemblies, which were then thoroughly quality controlled. All assemblies were performed with the relevant genome size parameter set to, or coverage calculation based on, a 1.2 Gbp genome size.

For coverage curves, pass reads were subset randomly to yield 40, 60, 80, and 100% of reads in each library. Canu version 1.3 + (commit: 37b9b80) was used for initial read correction with the parameters corOutCoverage=500, corMinCoverage=2, and minReadLength=2000 (later used as input for SMARTdenovo). Final Canu assemblies were performed with updated Canu version 1.4 + (commit: 0c206c9) and default parameters. Minimap^18^ (vesion 0.2-r124-dirty) was used to find overlaps with -L 1000 -m0 -Sw5, and miniasm (version 0.2-r137-dirty) was used to complete the assembly. For selected ‘top’ assemblies, miniasm and Canu were ran as above.

We tested several datasets as input to SMARTdenovo 61cf13d to compare the contiguity metrics of the resulting assemblies (Supplementary File D). The random subsets of reads (Subset040, Subset060, Subset080 and Subset100) were used but we also selected 30X of the longest raw reads and Canu-corrected reads (Supplementary File D, I). As it was previously demonstrated^19^, giving only a subset of the longest reads to SMARTdenovo could be beneficial to the assembly results. The assembler parameters were ‘-c 1’ to run the consensus step and ‘-k 17’, as a larger k-mer size than 16 is advised on large genomes with k=17. Wtdbg version 3155039 was ran with S=1.02, k=17. SMARTdenovo was ran on 30X coverage of the longest pass reads with k=17. The 30X coverage of the longest corrected reads was then assembled with SMARTdenovo using k=17.

Finally, parameters for additional miniasm and SMARTdenovo assemblies are detailed in (Supplementary File C,D), respectively.

#### BUSCO

Quality of genomes for gene detection was assessed with BUSCO (version 2.0)^22^ against the embryophyta_odb9 lineage. BUSCO in turn used Augustus (version 3.2.1)^25^, NCBI’s BLAST (version 2.2.31+)^26^, and HMMER (version 3.1b2)^27^.

#### Illumina read trimming

Illumina reads were trimmed for low quality bases and TruSeq-3 adapter sequences using Trimmomatic 0.35^28^ with a sliding window of 4-bases and average quality score threshold of 15. Reads below a minimal length of 36 base pairs after trimming were dropped.

#### k-mer analysis

A total of 25 billion 17-mers were generated from the adapter trimmed Illumina paired-end data using Jellyfish (v2.2.4)^29^. 17-mers with a depth of below 8 were considered error-prone and dropped for further analysis. The remaining 24 billion 17-mers indicated a peak depth of 22 resulting in a genome size estimate of 1.12 Gbp.

#### Polishing

##### Racon

Racon^23^ was used in version 0.5.0 based on overlaps created with the included minimap release. Both tools were used with standard settings except switching off the read quality filtering option in racon (--bq -1).

##### Pilon

Iterative polishing by Pilon (v1.20)^24^ was achieved by aligning adapter-trimmed paired-end Illumina reads to the corresponding assembly or polished consensus sequence from the previous iteration using bwa mem (v0.7.15-r1140)^30^. The resulting sorted alignment file (samtools v1.3)^21^ was subjected to Pilon^24^ (v1.20) together with the corresponding assembly for generation of a new consensus sequence. Pilon was run at default settings to fix bases, fill gaps and local misassemblies.

##### Qualimap

Illumina reads were mapped to the assemblies with bwa mem, secondary alignments were removed with samtools, and discrepancies were quantified with Qualimap (v.2.2.1).

##### Read Quality

Expected error rate was quantified across reads and pass / fail subsets of libraries according to the Phred scores in FASTQ files. Specifically: the sum of 10^phred / -10^ at each base position, divided by the number of bases. Empirical read quality was gathered by aligning nanopore reads back to the 4-times Pilon polished Canu assembly using bwa mem -x ont2d (v0.7.15-r1140) and calculating read identity including InDels per mapped bases.

##### Determination of summary statistics

Assembly statistics were computed using quast^31^ (v4.3) for eukaryotes (-e). Oxford Nanopore metadata and fastq sequences were extracted from basecalled fast5 files using in-house scripts.

##### Dotplots

Dotplots were generated using the MUMMER package^32^ (v.3.23). The unpolished assemblies were aligned to the reference genome of *S. pennellii* LA716^5^ using nucmer. The resulting alignment was filtered for a minimal alignment length of 20 kb (-l 20000) and 1-to-1 global alignments (-g) and subsequently partitioned based on chromosome. Plotting was performed using mummerplot.

## Gas Chromatography-Mass spectrometry (GC-MS)

Extraction and analysis by gas chromatography mass spectrometry was performed using the same equipment set up and exact same protocol as described in Lisec et al^33^. Briefly, frozen ground material was homogenized in 700 μL of methanol at 70°C for 15 min and 375 μL of chloroform followed by 750 μL of water were added. The polar fraction was dried under vacuum, and the residue was derivatized for 120 min at 37°C (in 60 μl of 30 mg ml-1 methoxyamine hydrochloride in pyridine) followed by a 30 min treatment at 37°C with 120 μl of MSTFA. An autosampler Gerstel Multi Purpose system was used to inject the samples to a chromatograph coupled to a time-of-flight mass spectrometer (GC-MS) system (Leco Pegasus HT TOF-MS). Helium was used as carrier gas at a constant flow rate of 2 ml/s and gas chromatography was performed on a 30 m DB-35 column. The injection temperature was 230°C and the transfer line and ion source were set to 250°C. The initial temperature of the oven (85°C) increased at a rate of 15°C/min up to a final temperature of 360°C. After a solvent delay of 180 sec mass spectra were recorded at 20 scans s-1 with m/z 70-600 scanning range. Chromatograms and mass spectra were evaluated by using Chroma TOF 4.5 (Leco) and TagFinder 4.2 software.

## Data availability

Data is available at http://www.plabipd.de/portal/solanum-pennellii. In addition data has been deposited at the EBI under accession PRJEB19787

## Acknowledgements

We want to acknowledge partial funding through the German Ministry of Education and Research 0315961 and 031A053 and 031A536C, the Ministry of Innovation, Science and Research within the framework of the NRW Strategieprojekt BioSC (no. 313/323-400-002 13), the DFG grant nos US98/7-1 and FE552/29-1 within ERACAPS Regulatome, and France Génomique (ANR-10-INBS-09).

## Author Contributions

B.U. designed the project. B.U. and A.M.B managed the project. M.H-W. S, A.V. and A.W developed the DNA extraction and sequencing protocol and generated primary sequencing data. J.M., M.E.B., A.M.B. processed and analyzed primary data. A.V., A.D., A.M.B., B.I, H.V., J-M. A., B.U. and A.R.F. conducted secondary data analyses, assemblies and statistics. S.A., A.R.F., D.Z., M.-H. W, C.P., U.S., R.C. analyzed plants, provided materials. F.M. provided material. A.D., M.H.-W., A.V., B.I., J.-M. A., A.M.B., D.Z., U.S., S.A., A.R.F, and R.C. interpreted data, and B.U. wrote the manuscript with help from all authors.

The authors declare no competing interests.

